# Structural Analysis and Docking Studies of FK506-Binding Protein 1A

**DOI:** 10.1101/2025.05.22.655516

**Authors:** Omur Guven, Hasan DeMirci

**Affiliations:** Koc University; SLAC National Laboratory

**Keywords:** FKBP1A, structural biology, X-ray crystallography, molecular docking

## Abstract

FK-binding protein 1A, a member of immunophilin family of proteins, is a protein with a wide variety of roles in cellular processes, including regulation of immune system, calcium intake metabolism through ryanodine receptors, TGF-β signaling and EGFR regulation. As a protein originally defined as the cellular target of premier immunosuppresant drugs, FK506 and Rapamycin, it has been a protein studied for further pharmacological uses. In this study, we have overexpressed, purified and crystallized apo-FKBP1A. Here, we are showing the FKBP1A crystal structure, calculated at cryogenic temperature at a very high resolution of 1.05 Å, obtained with a home source X-ray ‘Turkish DeLight’. Docking studies, with drug repurposing in mind were carried out with Molegro Virtual Docker software. Docking results will prove useful in future pharmaceutical studies done on FKBP1A, and similar proteins.

## 1. INTRODUCTION

FKBP1A (FKBP12), a member of the FK506-binding protein family, plays critical roles in cellular functions, including immunosuppression, calcium signaling, and protein regulation through proline cis-trans isomerization. FKBP proteins catalyze the interconversion of proline’s peptide bonds between cis and trans conformations, which profoundly influences protein structure and function. Among its diverse functions, FKBP1A interacts with immunosuppressive drugs FK506 and rapamycin, eliciting distinct cellular responses. FK506 inhibits calcineurin, preventing NFAT-mediated T cell activation, while rapamycin inhibits mTORC1, affecting cell growth and memory T cell differentiation **[Tong et al, 2015; Kasahara et al, 2021].** These mechanisms underline FKBP1A’s importance in immune modulation.

FKBP1A’s regulatory roles extend beyond immunosuppression. It stabilizes ryanodine receptors (RyRs), which regulate calcium release from the endoplasmic and sarcoplasmic reticulum, essential for cardiac excitation-contraction coupling. Dysregulation of this interaction may lead to arrhythmias or heart failure **[Gonano et al, 2017].** Additionally, FKBP1A interacts with Transforming Growth Factor-β (TGF-β) receptors, particularly regulating activin-receptor-like kinase (ALK) in the bone morphogenetic protein (BMP) pathway **[Stockwell et al, 1998; Quist-Løkken et al, 2021].** This role extends its relevance to autophagy and cancer, where FKBP1A acts as a biomarker and influences the ubiquitination and degradation of MDM2, a regulator of the tumor suppressor p53 **[Li et al, 2022: Cai et al, 2022].** Furthermore, FKBP1A modulates the autophosphorylation of the Epidermal Growth Factor Receptor (EGFR), highlighting its multifaceted regulatory capacity **[Mathea et al, 2011].**

In neurodegenerative diseases, FKBP1A’s role has garnered significant attention. Lower FKBP1A levels have been observed in Alzheimer’s disease (AD) brains, while elevated levels are found in neurofibrillary tangles and other pathologies**[Caminati et al, 2020; Chattopadhaya et al, 2011].** FKBP1A interacts with α-synuclein, accelerating its aggregation into fibrils associated with Parkinson’s disease (PD). This interaction exacerbates neurodegenerative processes, suggesting that FKBP1A inhibition could have therapeutic potential. Moreover, FKBP1A may regulate tau phosphorylation, contributing to aberrant protein folding and transport impairments characteristic of AD. Its interaction with calcineurin further implicates FKBP1A in calcium signaling dysregulation, a hallmark of neurodegeneration **[Avramut et al, 2002]**. Innovative detection methods, including nanostructured platforms, offer promise for FKBP1A’s use as a biomarker in neurodegenerative disease screening and therapeutic development.

FKBP1A also plays a pivotal role in cancer development and therapy. It modulates key signaling pathways, including TGF-β, apoptosis, and angiogenesis. For example, FKBP1A’s interaction with the TGF-β receptor suppresses tumor-promoting signaling under normal conditions. Loss of FKBP1A’s regulatory function has been linked to poor prognosis in cancers such as breast cancer and liver hepatocellular carcinoma (LIHC) **[Romano et al, 2015; Solassol et al, 2011].** High FKBP1A expression in LIHC correlates with advanced tumor stages and poor survival rates, while its loss in breast cancer predicts resistance to anthracycline-based therapies **[Theuerkorn et al, 2011; Romano et al, 2010].** Additionally, FKBP1A’s role in regulating HIF-2α and EGFR underscores its contribution to tumorigenesis through angiogenesis and apoptosis resistance **[Yao et al, 2011; Xing et al, 2019].** Emerging research suggests that FKBP family members, including FKBP1A, are valuable biomarkers and therapeutic targets across cancer types.

Structurally, FKBP1A’s functions are closely tied to its ability to bind ligands and regulate protein interactions. High-resolution studies of FKBP1A at cryogenic temperatures provide critical insights into its regulatory mechanisms **[Li et al, 2022].** Understanding these structural details advances its application in therapeutic contexts, whether targeting immune pathways, neurodegenerative diseases.

Overall, FKBP1A is a versatile protein with extensive roles in cellular regulation, disease progression, and therapeutic potential. Its involvement in diverse signaling pathways makes it a crucial target for future research aimed at understanding and manipulating its functions for clinical benefit.

## 2 METHODS

### 2.1 Transformation and Expression

FKBP1A with the sequence “GVQVETISPGDGRTFPKRGQTCVVHYTGMLEDGKK FDSSRDRNKPFKFMLGKQEVIRGWEEGVAQMSVGQRAKLTISPDYAYGATGHPGIIPPHATLVFDVELLKLE” was cloned into pET28a (+) vector with an N-terminal hexahistidine purification tag with a Thrombin cut site. As a cloning restriction enzyme cut sites, HindIII and KpnI were chosen and the Kanamycin resistance gene was used as a selection marker. The constructed plasmid was transformed into competent Escherichia coli (E. coli) BL21 Rosetta-2 strain, with heat shock method. Transformed bacterial cells were grown in 18 L regular LB media containing 50 µg/mL kanamycin and 35 µL/mL chloramphenicol at 37 °C. At OD600 value of 0.8, the protein expression was induced by using β-D-1-thiogalactopyranoside (IPTG) at a final concentration of 0.4 mM for 18 h at 18 °C. Cell harvesting was done using Beckman Allegra 15 R desktop centrifuge at 4 °C at 3500 rpm for 45 min. Cell pellets were stored at −45°C until protein purification.

### 2.2 Purification

The cells were dissolved in lysis buffer containing 500 mM NaCl, 50 mM Tris-HCl pH 7.5, 10% (v/v) Glycerol, 0.1% (v/v) Triton X-100, 2 mM BME, and 10 µM ZnCl2. The homogenized cells were lysed using a Branson W250 sonifier (Brookfield, CT, USA). The cell lysate was centrifuged at 4 °C at 35000 rpm for 1 h with Beckman Optima™ L-80XP Ultracentrifuge equipped with Ti45 rotor (Beckman, Brea, CA, USA). The pellet containing membranes and cell debris was discarded. The supernatant containing the soluble protein was filtered through 0.2 micron hydrophilic membrane and loaded to a Ni-NTA column that was previously equilibrated with a wash buffer containing 250 mM NaCl, 20 mM Tris-HCl pH 7.5 and 20 mM Imidazole. Unbound proteins were discarded by washing the column using a wash buffer. Then, the target protein (FKBP1A) was eluted using an elution buffer containing 250 mM NaCl, 20 mM Tris-HCl pH 7.5, 250 mM Imidazole,. Then, the eluted FKBP1A protein was dialyzed in a dialysis membrane (3 kDa MWCO) against a buffer with the same composition as the wash buffer for 3 h at 4 °C to remove excess imidazole. Dialyzed FKBP1A protein was cut using Thrombin protease to remove the hexahistidine-tag overnight at 4 °C.

### 2.3 Crystallization

The crystallization screening of N-terminal hexahistidine cleaved FKBP1A was performed using the sitting-drop microbatch under oil method against ∼3000 commercially available sparse matrix crystallization screening conditions in a 1:1 volumetric ratio in 72-Terasaki plates (Greiner Bio-One, Kremsmünster, Austria) as described in Ertem 2021 et al. **[Ertem, 2022]**. The mixtures were covered with 16.6 µL 100% paraffin oil (Tekkim Kimya, Istanbul, Türkiye). The crystallization plates were incubated at 4 °C and checked frequently under a stereo light microscope. The best FKBP1A crystals were grown within three months in NR LBD condition #48 (Molecular Dimensions, USA). This condition contains 2.0 M Ammonium Sulfate, 0.2 M Sodium Chloride and 0.1M TRIS 8.0.

### 2.4 Crystal Harvesting and Delivery

The FKBP1A crystals were harvested using MiTeGen MicroLoops attached to a magnetic wand **[Garman, 2006]** while being monitored under microscope **[Atalay, 2023]**. The obtained crystals were flash frozen by plunging in liquid nitrogen and placed in a cryo-cooled sample storage puck (Cat#M-CP-111-021, MiTeGen, USA). Then, the puck was placed into the Turkish DeLight liquid nitrogen-filled autosample dewar at 100 K.

### 2.5 Data Collection and Data Reduction

Collection of diffraction data from the FKBP1A crystal was performed by utilizing Rigaku’s XtaLAB Synergy Flow XRD source “*Turkish DeLight*” at University of Health Sciences (Istanbul, Türkiye) with *CrysAlisPro* software 1.171.42.35a **[22]**. The crystals were kept cooled by the Cryostream 800 Plus system, which was set to 100 K. The PhotonJet-R X-ray generator operated at 30 mA, 1200.0 W, and 40 kV with 10% beam intensity. The data was collected at 1.54 Å wavelength and the detector distance was set to 47.00 mm. The crystal oscillation width was set to 0.25 degrees per image while exposure time was 20.0 min. CrysAlis Pro version 1.171.42.35a **[22]** was utilized to perform data reduction and an *.mtz file was obtained as the result.

### 2.6 Structure Determination and Refinement

The cryogenic TRAF6 structure was determined at 1.05 Å with the space group C121 utilizing *PHASER* 2.8.3 **[McCoy, 2007],** an automated molecular replacement program within the *PHENIX* suite 1.20.1 **[Adams, 2010].** An AlphaFold generated FKBP1A structure was used as an initial search model for molecular replacement. Simulated annealing and rigid-body refinements were performed during the first refinement cycle including individual coordinates and translation/liberates/screw (TLS) parameters were refined. The structure was checked by *COOT* **[Emsley, 2004]** after each set of refinement and the solvent molecules were added into unfilled, appropriate electron density maps. The obtained final structure was examined by using *PyMOL* **[26]** 2.5.4 and *COOT* 0.9.8, and the figures were created. Data collection and refinement statistics were given in Table 1.

**Table 1.** Data collection and refinement statistics.

|  | **FKBP1A** |
| --- | --- |
| **Wavelength** |  |
| **Resolution range** | 22.08 - 1.05 (1.088 - 1.05) |
| **Space group** | C 1 2 1 |
| **Unit cell** | 101.85 36.124 54.499 90 95.987 90 |
| **Total reflections** | 405957 (19637) |
| **Unique reflections** | 90348 (6413) |
| **Multiplicity** | 4.5 (2.2) |
| **Completeness (%)** | 87.38 (69.72) |
| **Mean I/sigma(I)** | 10.74 (0.65) |
| **Wilson B-factor** | 9.46 |
| **R-merge** | 0.07648 (1.222) |
| **R-meas** | 0.0824 (1.545) |
| **R-pim** | 0.02891 (0.9284) |
| **CC1/2** | 0.999 (0.231) |
| **CC\*** | 1 (0.613) |
| **Reflections used in refinement** | 80574 (6414) |
| **Reflections used for R-free** | 2000 (159) |
| **R-work** | 0.2050 (0.4140) |
| **R-free** | 0.2235 (0.4278) |
| **CC(work)** | 0.971 (0.582) |
| **CC(free)** | 0.965 (0.592) |
| **Number of non-hydrogen atoms** | 2109 |
| **macromolecules** | 1773 |
| **ligands** | 10 |
| **solvent** | 326 |
| **Protein residues** | 216 |
| **RMS(bonds)** | 0.008 |
| **RMS(angles)** | 1.13 |
| **Ramachandran favored (%)** | 97.17 |
| **Ramachandran allowed (%)** | 2.83 |
| **Ramachandran outliers (%)** | 0.00 |
| **Rotamer outliers (%)** | 1.04 |
| **Clashscore** | 2.80 |
| **Average B-factor** | 14.25 |
| **macromolecules** | 12.84 |
| **ligands** | 8.90 |
| **solvent** | 22.10 |
| **Number of TLS groups** | 14 |

Statistics for the highest-resolution shell are shown in parentheses.

### 2.7 Molecular Docking

After Traf6 structure was determined with home source, at cryogenic temperature, molecular docking was done by utilizing Molegro Virtual Docker(MVD) **[Bittencout-Ferreira, 2019].** For this step, entire FDA database was obtained from Drugbank server **[Knox, 2024].** Then, the active sites on Traf6 were determined using MVD and the docking procedure was followed.

## 3 RESULTS

### 3.1 FKBP1A Structure at 1.05 A

FKBP1A is a 24 kDa dimerized protein and here we present a cryogenic temperature full length structure at 1.05 Å. (Figure) The presented structure is also deposited to Protein Database (PDB) with the accession code 8X6P.

**Figure 1:**
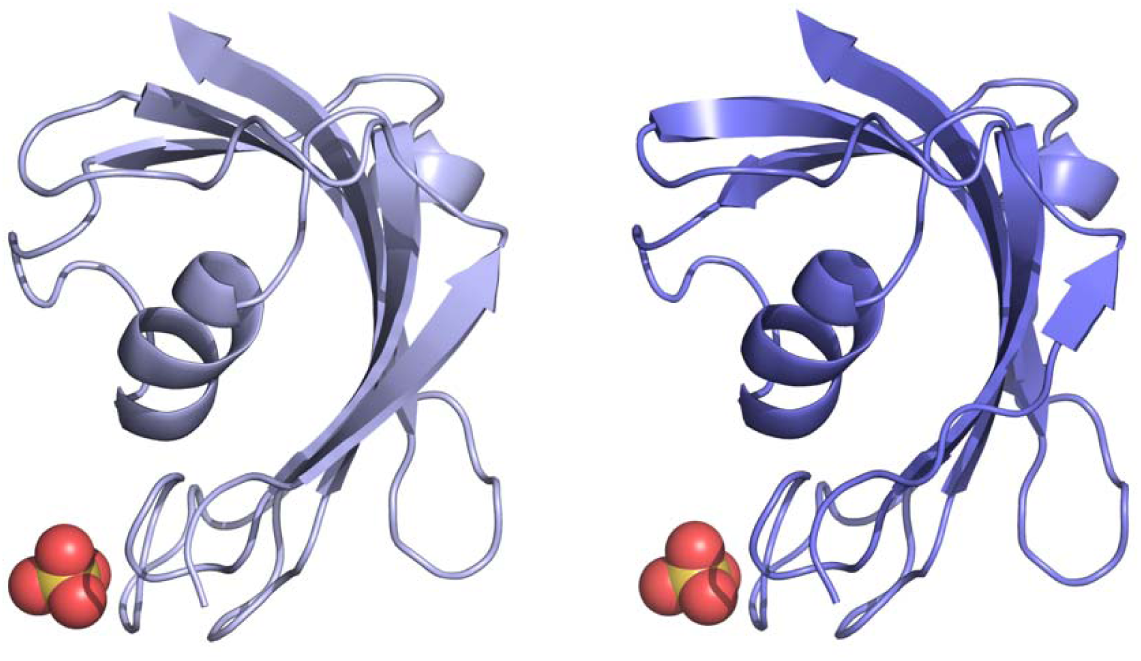
Overall structure of FKBP1A with sulfate ions present. These sulfate ions come from the crystallization condition (NR LBD-II, #48) which includes Ammonium Sulfate.

As we look into the sulfate ions in detail, we can see that they are interacting with Ala85 and Thr86, located in the loop region.

**Figure 2:**
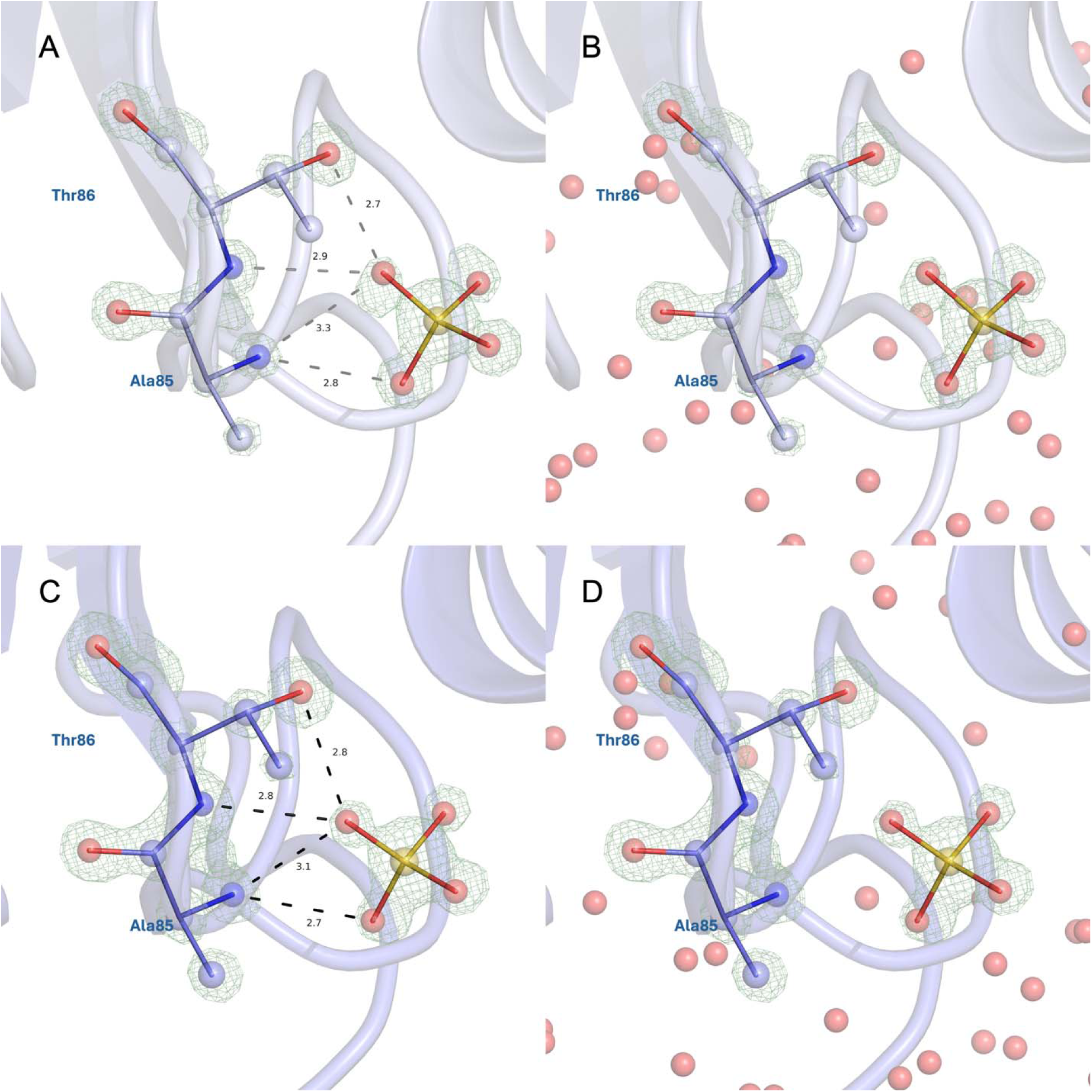
Sulfate interaction with Ala85 and Thr86. **(A,C)** Distances between SO4 and interacting ions in Chain A and Chain B, respectively, shown with electron density maps at sigma level of 4.0 **(B,D)** Electron density maps showing the sulfate and the interacting residues in Chain A and Chain B, respectively, with the surrounding solvent residues.

### 3.2 Structural Alignment with Reference Model

The AlphaFold model used for molecular replacement was aligned with the determined FKBP1A structure. As the model is a monomeric structure, we have aligned with Chain A of our structure here. Two structures align with a RMSD value of 0.263, with the main diffraction being the sulfate ion in our structure.

**Figure 3:**
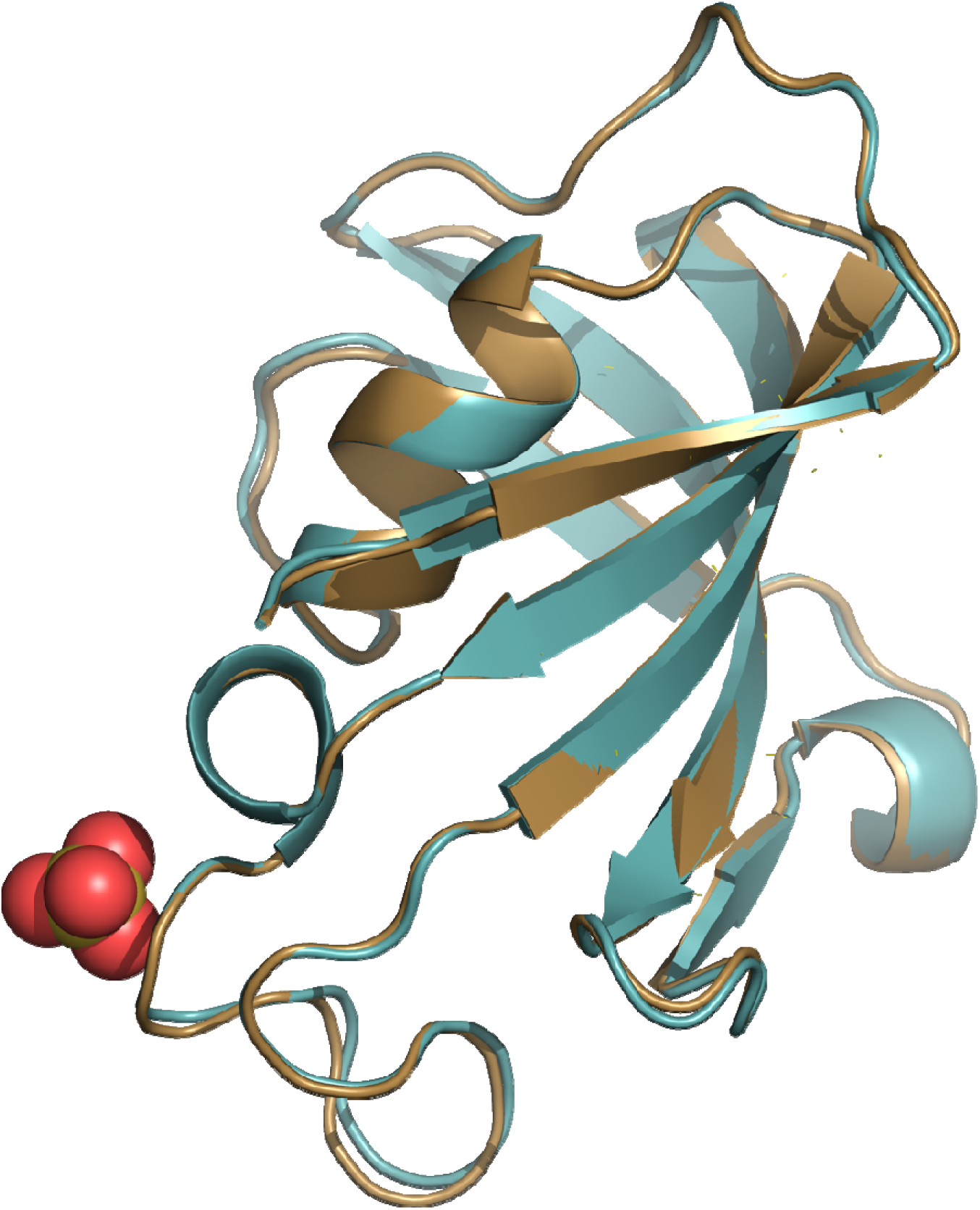
FKBP1A Chain A aligned to AlphaFold model with RMSD 0.263. Our structure has a sulfate ion, while the model structure does not possess one.

### Molecular Docking for FKBP1A

The FDA database, obtained from Drugbank, was screened on the full length FKBP1A (PDB accession code: 8X6P) by using Molegro Virtual Docker (MVD). For this analysis, the possible binding pockets were determined first, and the pocket between the two chains were selected as the target. Then, 3353 compounds were screened in this pocket, and were ranked based on their ‘Rerank score’, a function of MVD. Shared in Table 2 is the Rerank scores and molecular weights for top 10 candidates.

**Table 2:**
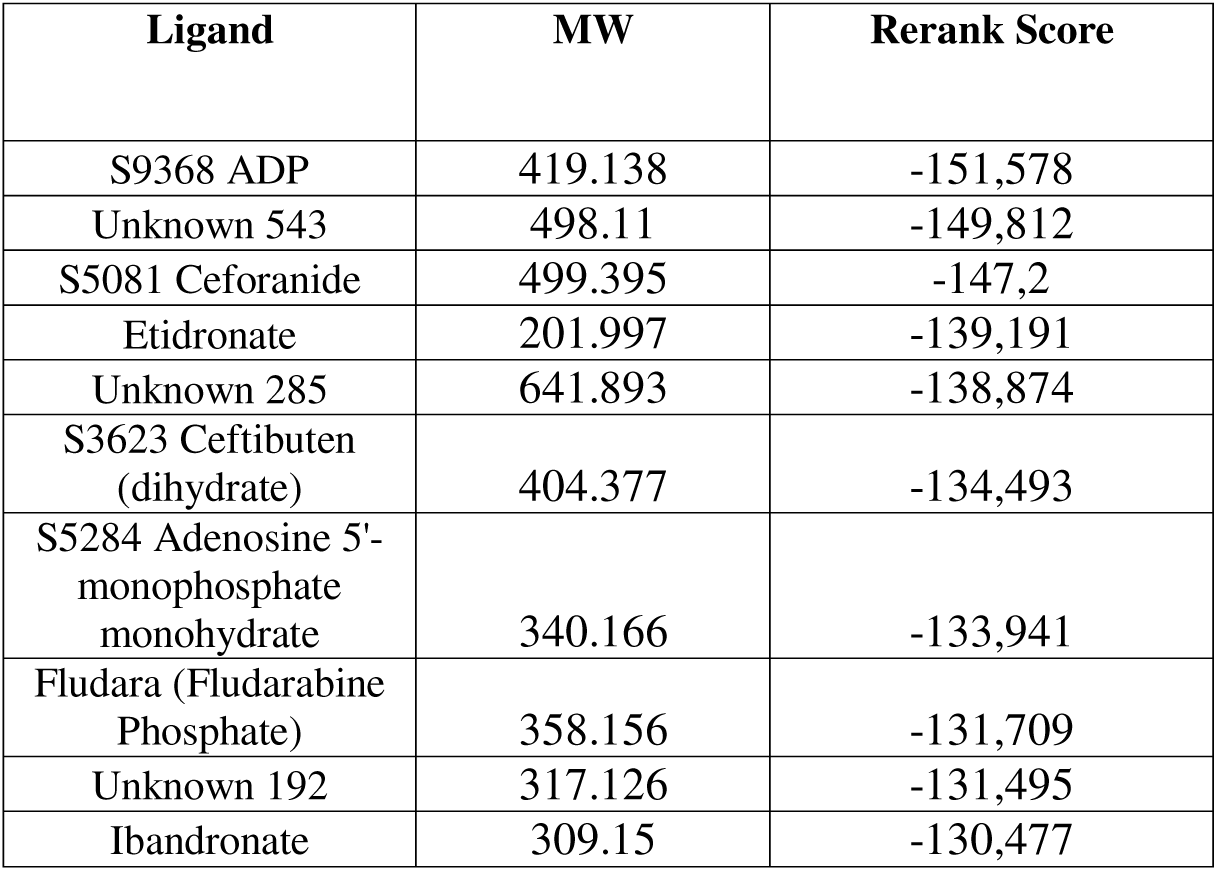
Rerank scores and molecular weights of the top 10 candidates of molecular docking.

These rerank scores are calculated as the summation of different binding free energies of intermolecular interactions. These interactions can be categorized as protein-ligand, cofactor-ligand and solvent(water)-ligand. In this case, the cofactor is the sulfate ion, shown in the structure. These values are shown in Table 3.

**Table 3:** Intermolecular energy calculations for top 10 compounds.

| Ligand | E_inter<br>(protein_ligand) | E_inter<br>(cofactor_ligand) | E_inter<br>(water_ligand) |
| --- | --- | --- | --- |
| S9368 ADP | -135,746 | -26.7246 | -35.5842 |
| Unknown 543 | -127,822 | -36.8172 | -48.4087 |
| S5081 Ceforanide | -108,373 | -17.8779 | -51.4766 |
| Etidronate | -108,528 | -39.8666 | -26.0207 |
| Unknown 285 | -134,389 | -91.446 | -38.9176 |
| S3623 Ceftibuten<br>(dihydrate) | -106,103 | -17.5336 | -41.1327 |
| S5284 Adenosine 5'-<br>monophosphate<br>monohydrate | -113,662 | -17.9274 | -34.0511 |
| Fludara (Fludarabine<br>Phosphate) | -115,586 | -18.085 | -30.9713 |
| Unknown 192 | -113,594 | -17.9217 | -29.577 |
| Ibandronate | -122,311 | -38.6525 | -28.1673 |

Docking studies also show the interactions ‘opposing’ the pose, such as clashes. These values could be important, as they can arise from an unfavorable pose for the compound. It should be noted that these values are related to the conformation of the compound only, instead of being an indication of how well they are bound to the protein. Intramolecular energy values are shown in Table 4.

**Table 4:** Intramolecular energy calculations for top 10 compounds.

| Ligand | E_intra (steric) | E_inter (tors) | E_intra (vdW) |
| --- | --- | --- | --- |
| S9368 ADP | 19.4346 | 6.93365 | 33,72 |
| Unknown 543 | 33.6202 | 16.028 | 57,1525 |
| S5081 Ceforanide | -10.6038 | 0 | 72,9067 |
| Etidronate | 5.20588 | 0 | -2,1155 |
| Unknown 285 | 41.028 | 18.4005 | 224,859 |
| S3623 Ceftibuten<br>(dihydrate) | 9.79644 | 0 | 50,6147 |
| S5284 Adenosine 5'-<br>monophosphate<br>monohydrate | 4.55114 | 2.83817 | 62,3046 |
| Fludara (Fludarabine<br>Phosphate) | -0.10244 | 4.30578 | 46,8188 |
| Unknown 192 | 13.3931 | 2.82438 | 34,7208 |
| Ibandronate | 16.5892 | 0 | 110,434 |

### 3.3 Visualization and Ligand Interaction Maps of Top 10 Compounds

Once the top 10 cancidates are determined based on their rerank score, a more detailed examination was done on how they might interact with the receptor protein. For this purpose, the poses and the receptor protein (FKBP1A) were exported and were analyzed using Biovia Discovery Studio and PyMOL software suite.

**Figure 4:**
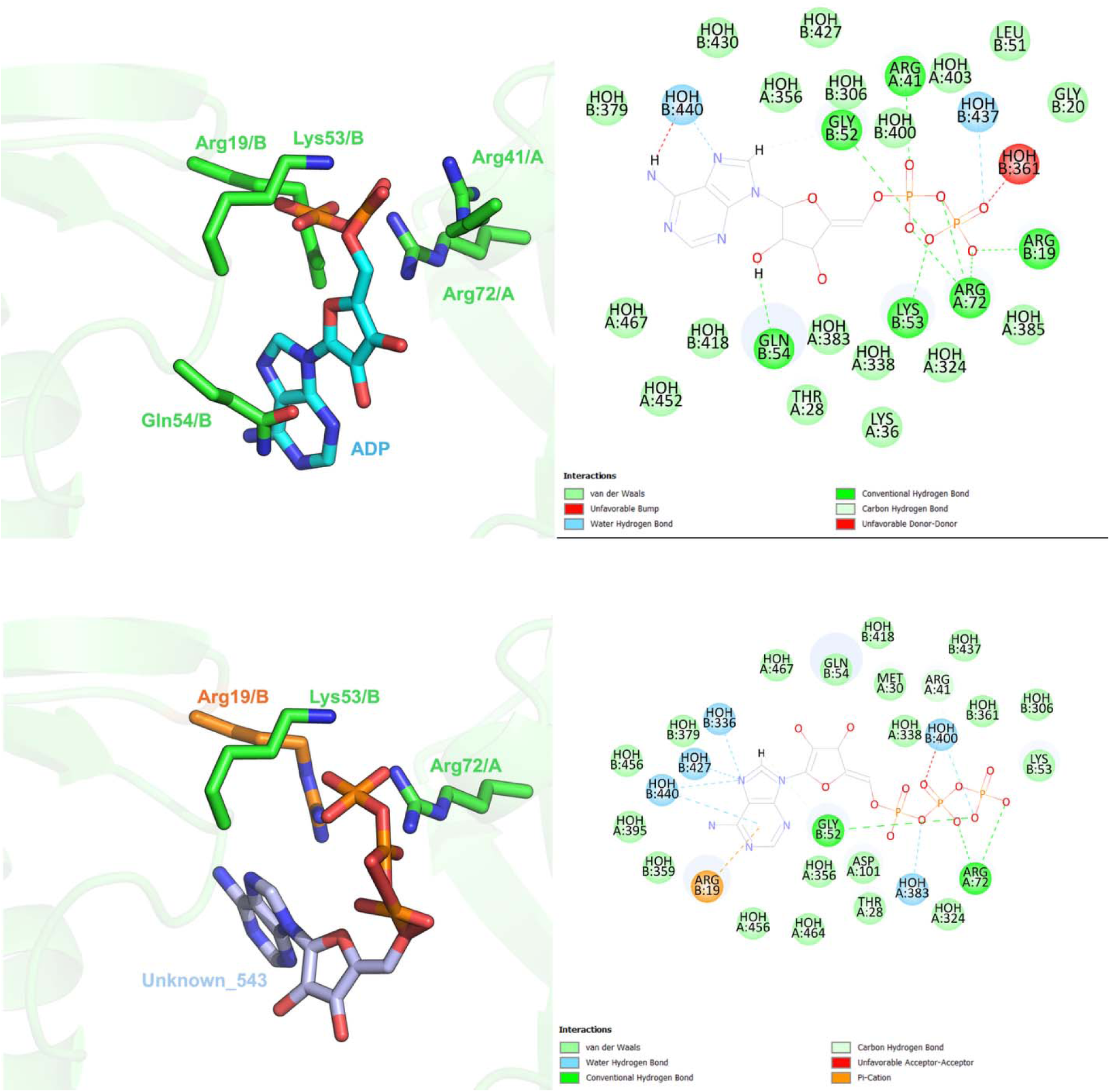
Proposed ADP and Unknown_543 bindings to FKBP1A and ligand interaction map. Residues from both chains are visualized here, and 2D interaction map shows hydrogen bonds, van der Waals interactions, interacting solvents and steric clashes.

**Figure 5:**
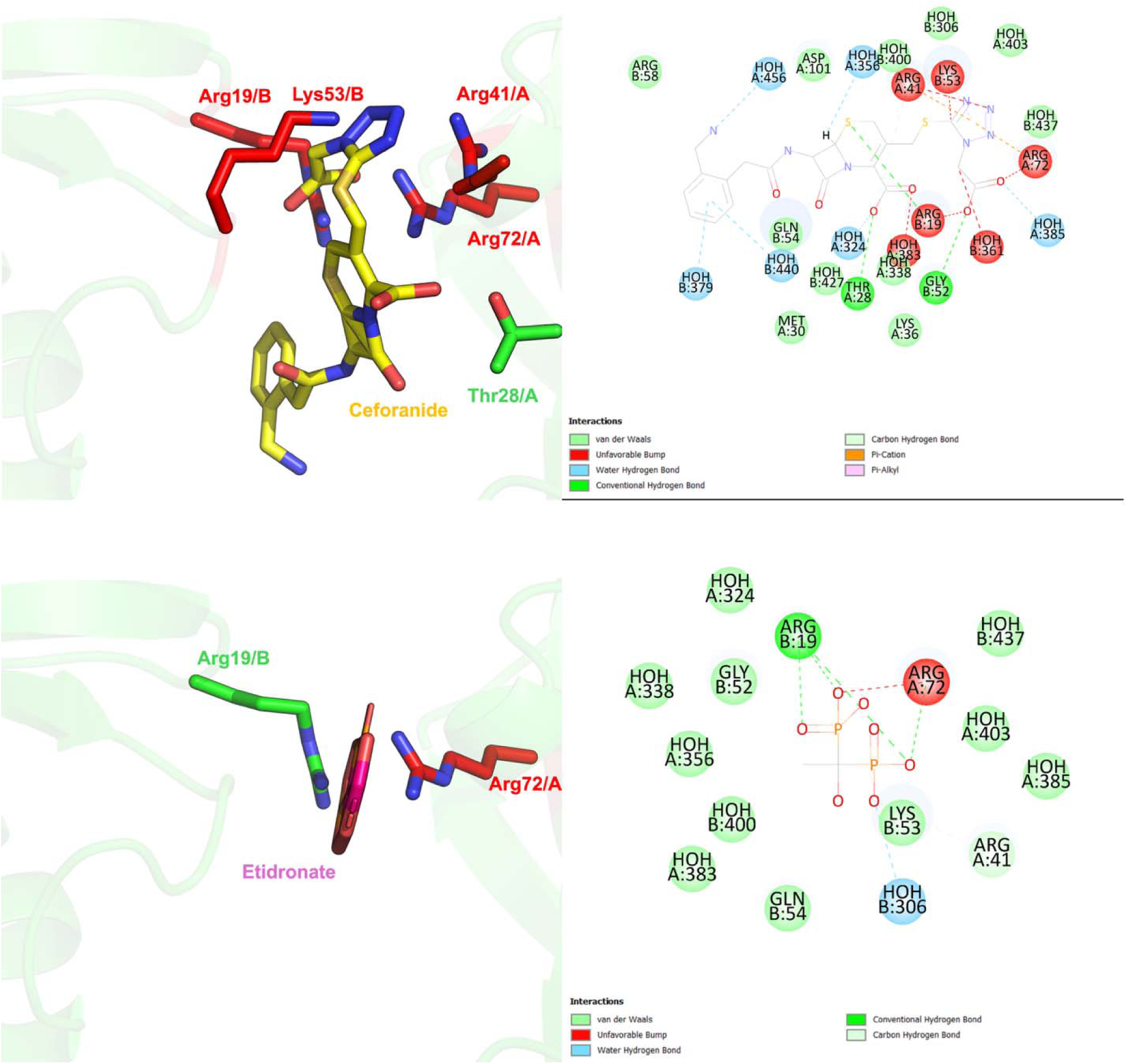
Proposed Ceforanide and Etidronate bindings to FKBP1A and ligand interaction map. Residues from both chains are visualized here, and 2D interaction map shows hydrogen bonds, van der Waals interactions, interacting solvents and steric clashes.

**Figure 6:**
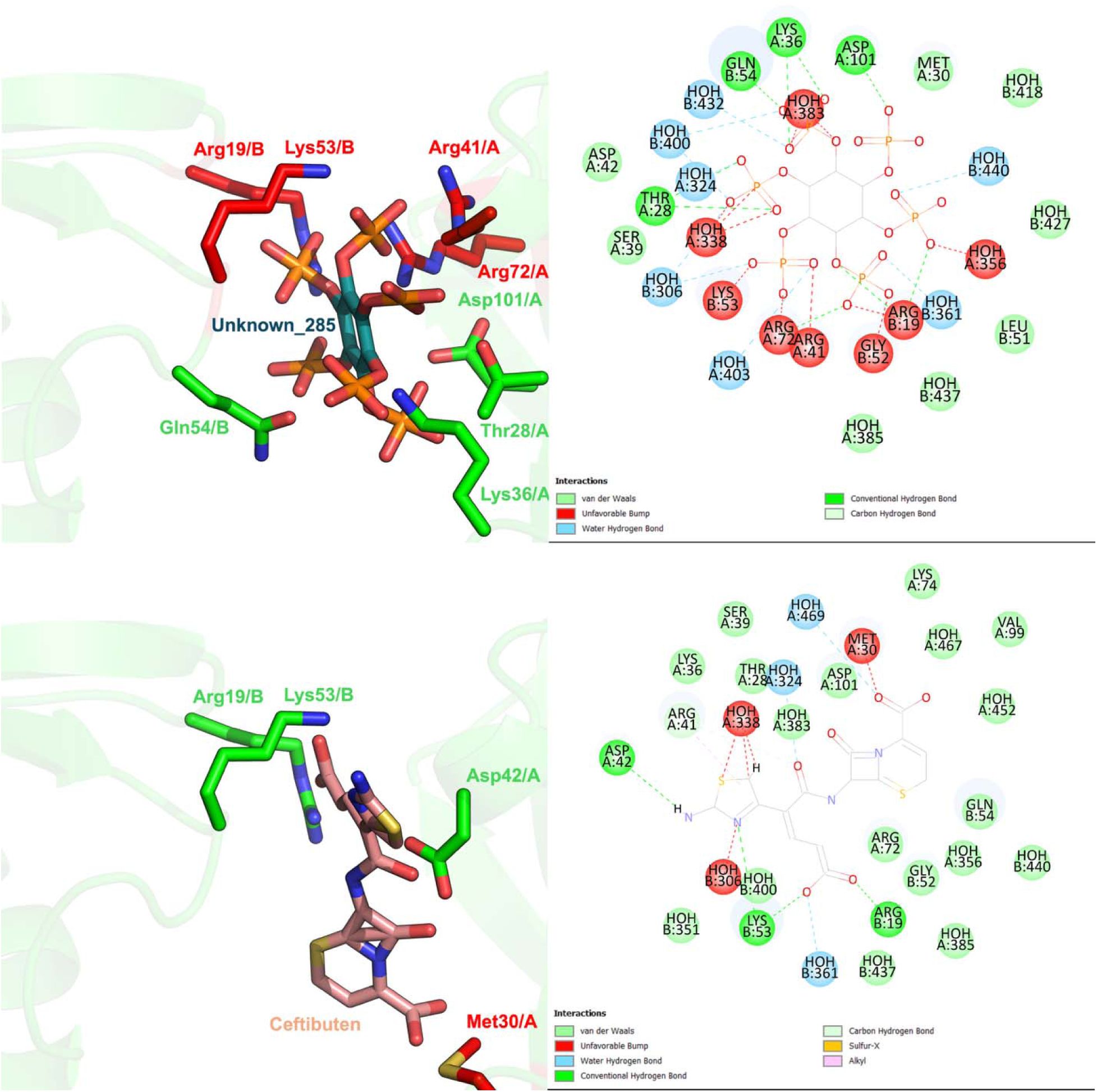
Proposed Unknown_285 and Ceftibuten bindings to FKBP1A and ligand interaction map. Residues from both chains are visualized here, and 2D interaction map shows hydrogen bonds, van der Waals interactions, interacting solvents and steric clashes.

**Figure 7:**
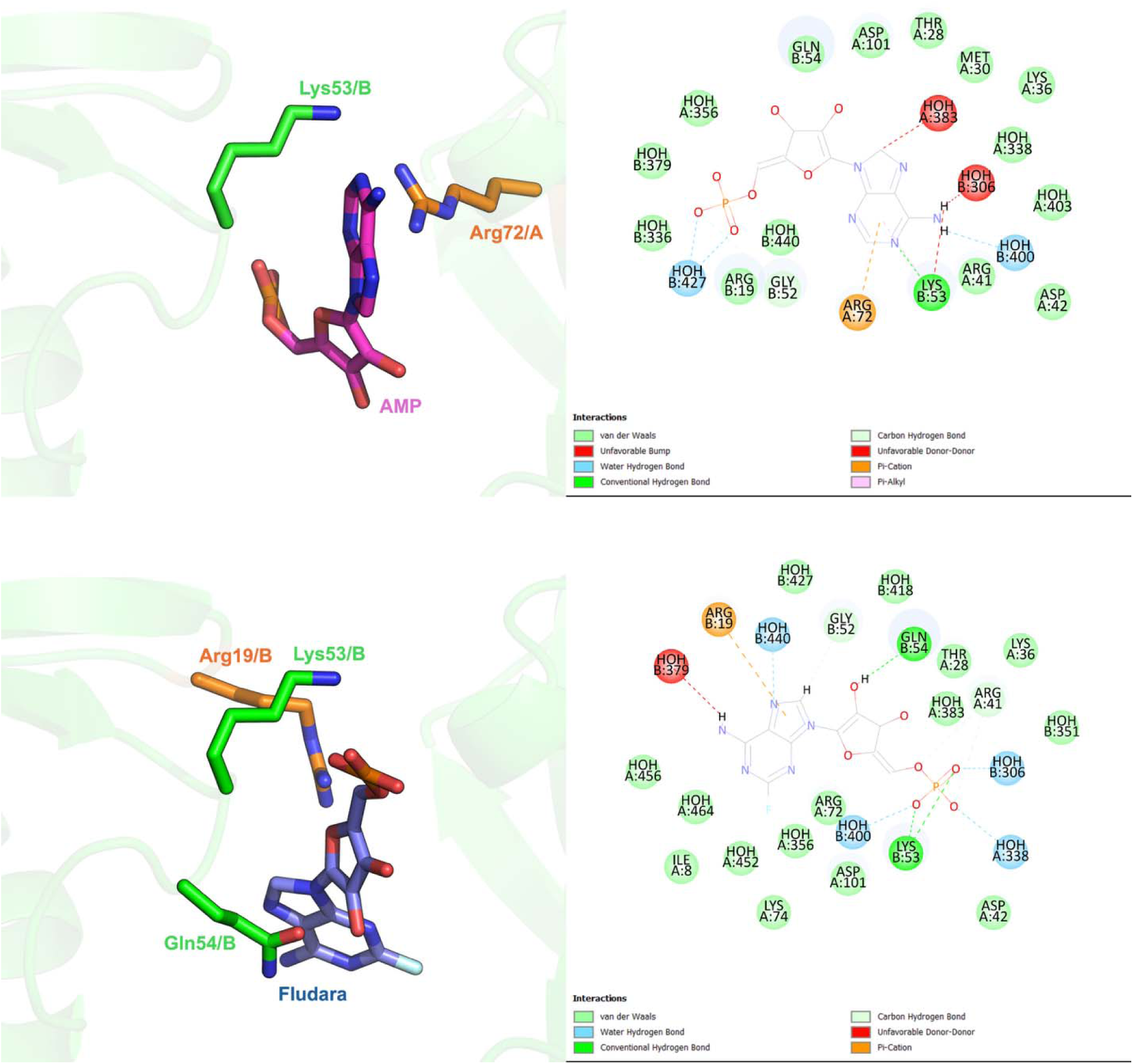
Proposed AMP and Fludara bindings to FKBP1A and ligand interaction map. Residues from both chains are visualized here, and 2D interaction map shows hydrogen bonds, van der Waals interactions, interacting solvents and steric clashes.

**Figure 8:**
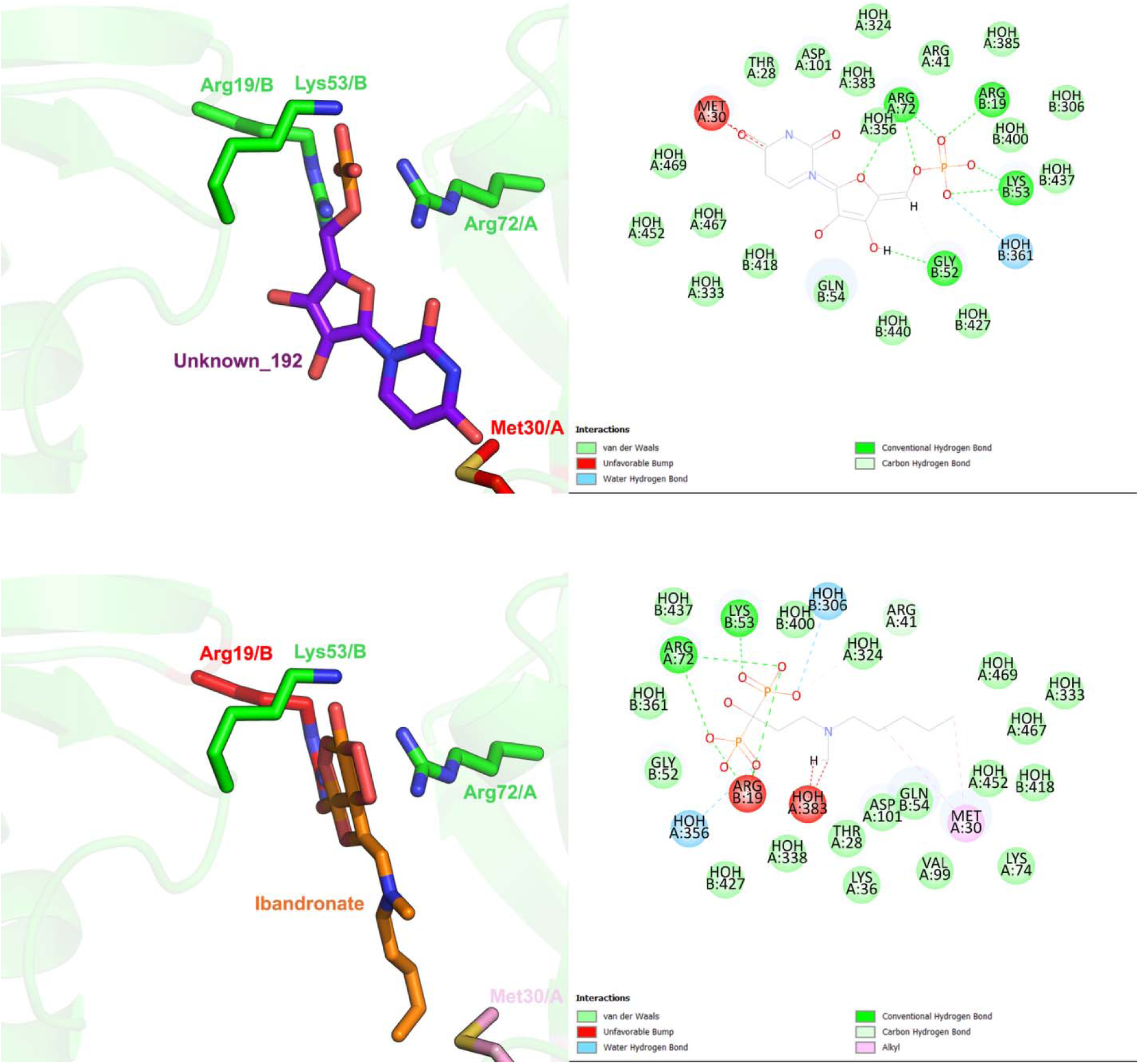
Proposed Unknown_192 and Ibandronate binding to FKBP1A and ligand interaction map. Residues from both chains are visualized here, and 2D interaction map shows hydrogen bonds, van der Waals interactions, interacting solvents and steric clashes.

## 4 DISCUSSION

FK506 Binding Protein (FKBP) family is a group of proteins with a wide array of roles, including immune system mediation, cell cycle regulation, neuronal signaling and neurodegeneration. Founding member of this family, FKBP1A, is a 12 kDa protein with a wide variety of cellular functions**[Hausch, 2015]** and a high conservation rate FKBP1A has a unique role as a peptidyl prolyl cis-trans isomerase, where it regulates the proline N-terminal peptide bond conformation. As the structure and the function of the protein is affected by this conformational change, FKBP1A is deemed to have a role in folding and regulation of especially proline rich proteins. It is also known for its role in stabilization of ryanodine receptors, preventing Ca^+2^ ions from endoplasmic reticulum.**[Gant, 2014]** In heart, altered levels of Ca^++^ can cause arrythmia and heart failure, thus demonstrating the importance of mediation. FKBP1A is also shown as a cellular interactor or TGF-ß Type I receptors, such as BMPs.

FKBP1A is also involved in various neurodegenerative diseases. In Alzheimer’s patients, FKBP1A is shown to colocalize in brain with neurofibrillatory tangles. It is also shown to interact with Amyloid Precursor Protein (APP). **[Cao, 2011]** In patients with Parkinson’s Disease, there is a connection between FKBP1A and α-Synuclein. **[Gerard, 2011]** It can also have a role in other tauopathies. **[Koren, 2011]** FKBP1A is thought to be involved in these diseases as it has a Peptidyl-prolyl cis-trans isomerase function.

In addition to its physiological roles, FKBP1A is a known target of common immunosuppressants, namely Rapamycin and FK506. These two drugs bind to the same protein, yet they trigger different cellular processes. Rapamycin binding to FKBP1A regulates mTOR pathway, a premier pathway involved in many cellular processes, such as cell cycle and cell survival. FK506 binding regulates calmodulin dependent calcineurin inhibition, which in turn affects the activity of NF-AT, a nuclear factor with a variety of roles in immune system regulation.**[Zissimopolus, 2021]**

Our FKBP1A model has two monomers in the asymmetric unit, and a pocket drawn between these two monomers during docking. The results show similar residues interacting with these compounds, such as a Lys53 residue in Chain B forming a hydrogen bond with 7 of them. On the other hand, the same residue causes a steric clash with two of the compounds. Last compound forms a van der Waals interaction with Lys53. Another such residue is Arg19 in Chain B, forming a hydrogen bond with four compounds, and a pi-cation interaction with 2 other. It causes a steric clash with 3 compounds, and a van der Waals interaction with the last one.

In Chain A, Arg 72 seems to be a key residue, forming a hydrogen bond with four compounds, and causing a clash with three others. It forms a van der Waals interaction with two compounds, and a pi-cation interaction with the last.

These three residues seem commonly interacting with the docked compounds, as illustrated here in ligand interaction maps. It is important to note that, this is the first reported FKBP1A structure with multiple chains and a sulfate ion, and the docking results also take these factors into account.

These findings and analyses could be important in studying the inhibition and uncovering new roles of FKBP1A in cellular processes.

## Acknowledgement

The authors gratefully acknowledge use of the services and facilities of the Koç University Isbank Infectious Disease Center (KUISCID), and Health Sciences University (SBÜ). H.D. acknowledges support from NSF Science and Technology Center grant NSF-1231306 (Biology with X-ray Lasers, BioXFEL). Ö.G. acknowledges support from the Scientific and Technological Research Council of Türkiye (TÜBİTAK, 2218 - National Postdoctoral Research Fellowship Program under project number 118C476). This publication has been produced benefiting from the 2232 International Fellowship for Outstanding Researchers Program and the 1001 Scientific and Technological Research Projects Funding Program of the TÜBİTAK (Project Nos. 118C270 and 120Z520). However, the entire responsibility of the publication belongs to the authors of the publication. The financial support received from TÜBİTAK does not mean that the content of the publication is approved in a scientific sense by TÜBİTAK.

